# Ultra-small Particle Size Nanobubbles Water Drinking Exerts Antifatigue Effects in Chronic Exercise Fatigue Rats via Improving Antioxidative Activities

**DOI:** 10.1101/2024.11.21.624464

**Authors:** Jie Qi, Tieyi Yang, Yinyan Zhao, Yinuo Liu, Maorui Han, Yue Liu, Jiayin Zhao, Jun Zhang

## Abstract

Chronic exercise fatigue is a prevalent issue that can lead to a decline in physical performance, metabolic dysregulation, and oxidative damage, posing significant health risks. This study aims to investigate the effects of Ultra-small Particle Size Nanobubbles Water (UsNBsW) which was Koishio technology produced water on physiological and biochemical parameters in a rat model of chronic exercise fatigue. We established a chronic fatigue model, and evaluated various physiological and biochemical indicators. Our key findings revealed that Koishio water significantly improved rats’ swimming performance, reduced blood lactate levels, and enhanced antioxidant capacity, as evidenced by increased hemoglobin content, decreased lactate dehydrogenase and creatine kinase activities (p<0.01). Additionally, liver glycogen levels were notably elevated (p<0.05). Futhermore, the antioxidant markers, including superoxide dismutase (SOD) and glutathione peroxidase (GSH-Px) activities were significantly higher (p<0.01), while malondialdehyde (MDA) and 8-hydroxydeoxyguanosine (8-OH-dG) levels were markedly reduced (p<0.01). Our results provide new clues for the promising role of Koishio water in mitigating chronic exercise fatigue and supply a reliable reference for future research into nutritional interventions and recovery strategies in exercise physiology.

## 1. Introduction

Chronic exercise fatigue is a prevalent issue that affects the physiological functions of various organisms, including humans and animal models. This syndrome can lead to a decline in physical performance, metabolic disturbances, and oxidative stress, which can ultimately compromise overall health. The importance of understanding chronic exercise fatigue lies in its implications for athletic performance, rehabilitation, and the management of certain medical conditions. Current research indicates that while several interventions have been explored to mitigate the effects of exercise-induced fatigue, there remains a significant gap in the systematic examination of specific dietary components, such as specialized waters, on chronic exercise fatigue in animal models [1,2].

Among the various interventions studied, the role of hydration and its constituents has been recognized as crucial for optimal physical performance and recovery. Prior studies have highlighted that hydration status can directly influence exercise capacity and recovery processes, with specific electrolytes and minerals being linked to improved performance outcomes. However, the existing literature lacks a comprehensive exploration of how unique water formulations, such as high-quality ultra-small particle size nanobubbles water, may specifically influence chronic exercise fatigue and the associated physiological responses in animal models [3]. This presents an opportunity for research aimed at elucidating the biochemical and physiological mechanisms underlying these effects.

The proposed research focuses on the evaluation of ultra-small particle size nanobubbles water (UsNBsW) and its effects on physiological and biochemical parameters in a chronic exercise fatigue rat model. This approach is informed by previous findings that suggest the composition of water can affect recovery and metabolic status following exercise. By systematically analyzing various physiological parameters, this study aims to fill a critical gap in the understanding of how specific water types can influence recovery from exercise fatigue, which has significant implications for the development of effective fatigue management strategies [4].

The methodology of this study includes establishing a rat model of chronic exercise fatigue, designing an appropriate exercise regimen, and assessing multiple physiological and biochemical markers. This comprehensive approach enables a detailed investigation of the specific impacts of UsNBsW on exercise-induced fatigue, thereby providing insights into potential recovery strategies that may benefit both athletic populations and those experiencing exercise-related fatigue in clinical settings [5].

The primary objective of this research is to elucidate the effects of UsNBsW on exercise performance, antioxidant capacity, and metabolic parameters in rats suffering from chronic exercise fatigue. By investigating these effects, the study aims to contribute valuable information that can inform future interventions and therapeutic strategies targeting exercise fatigue, ultimately enhancing performance and recovery in various populations [6].

In summary, this study addresses a significant research gap by investigating the influence of UsNBsW on chronic exercise fatigue in a rat model. The findings are anticipated to provide novel insights into the potential benefits of UsNBsW for improving recovery and performance, thereby advancing the field of sports science and clinical rehabilitation [7].

## 2. Materials and Methods

### 2.1 Animals

Fifty adult male Sprague-Dawley rats (200 ± 20g) were randomized into 5 groups: (1) Control group (CON group, n=10): normal feeding for 10 weeks, and drank boiled water daily; (2) Control + UsNBsW group (KW CF group, n=10): normal feeding for 10 weeks, and drank Ultra-small Particle Size Nanobubbles Water (UsNBsW) daily; (3) Chronic exercise fatigue group (Ex CF, n=10): rats continued to swim for 10 weeks of high intensity exercise, and drank boiled water; (4) Chronic exercise fatigue + UsNBsW group (KW-Ex CF, n=10): rats continued to swim for 10 weeks, and drank UsNBsW every day; (5) Chronic exercise fatigue + mineral water group (MW-Ex CF, n=10): rats continued to swim for 10 weeks, and drank mineral water every day. The UsNBsW were applied by Koishio technology produced water (KW). Following the BEDFORD animal exercise training load guidelines and based on the exercise training program outlined by FERNANDES et al., the rats in the exercise group swam in a 150 × 60 × 70 cm^3^ pool during a 10-week period. The water temperature was kept at 31 ± 1°C, the room temperature was maintained at 25 ± 2°C, air humidity was set at 50 ± 5%, and the light cycle was 12 hours per day. The protocol for this animal study adhered to the standards established in the Guide for the Care and Use of Laboratory Animals.

### 2.2 Exercise protocola

Rats were placed in a pool to perform swimming exercise without load. If a rat was submerged and did not resurface within 10 seconds, it was deemed exhausted. The exhausted rats were promptly taken out of the water and dried quickly. Some exhausted rats in a brief period were allowed to rest for 5 minutes before continuing to swim, aiming for a total exercise duration of 2 hours. Each rat swam once a day, engaging in exercise 5 days a week for a period of 10 weeks.

### 2.3 Sample collection

Ten weeks after the initiation of the study, the rats underwent testing for locomotor capabilities through a single exhaustive swimming trial. Following this swimming exercise, based on the body weight of the rats, they were administered an intraperitoneal injection of sodium pentobarbital at a concentration of 2% and a dosage of 1.25 g/kg for anesthesia. 1 mL of whole blood was collected from the abdominal aorta, and serum was separated with the addition of heparin for anticoagulation. The samples were then subjected to centrifugation at 2500 rpm for 15 minutes at room temperature to obtain the supernatant for analysis. Additionally, samples from the bilateral skeletal muscles, including the quadriceps, gastrocnemius, and soleus, were collected. After rinsing the remaining skeletal muscle tissue with normal saline, rapid immersion in liquid nitrogen was performed for subsequent testing.

### 2.4 Exercise endurance test

After ten weeks, the duration of swimming exercise was recorded by exhaustion exercise.

### 2.5 Determinations of Exercise fatigue index

Automatic blood analyzer for hemoglobin (Hb), blood lactic acid (LA), blood urea nitrogen (BUN), lactate dehydrogenase (LDH) and creatine kinase (CK) by the colorimetric method, and enzyme-linked immunoassay (Elisa) for serum testosterone (T) and corticosterone (C) determination.

### 2.6 Detection of blood glucose and glycogen levels

Blood glucose, muscle glycogen and liver glycogen levels were measured by ananthrine method.

### 2.7 Antioxidant index assay

The evaluation of rat serum and skeletal muscle catalase (CAT) activity, superoxide dismutase (SOD) activity, malondialdehyde (MDA) levels, and glutathione peroxidase (GSH-Px) activity was performed using a colorimetric method. Additionally, an ELISA technique was employed to measure the content of 8-hydroxydeoxyguanosine (8-OH-dG) in an animal model of chronic exercise fatigue.

### 2.8 Determinations of inflammatory index C-reactive protein

To detect C-reactive protein (CRP) content by elisa in rat serum and skeletal muscle in an animal model of chronic exercise fatigue.

### 2.9 Adenosine triphosphate assay

The adenosine triphosphate (ATP) levels in the serum and skeletal muscle of rats were measured using a colorimetric method in a model of chronic exercise-induced fatigue.

### 2.10 Statistical analysis

The data were expressed as mean ± SEM and were assessed using a one-way analysis of variance (ANOVA), with the Student-Newman-Keuls post hoc test applied, unless stated otherwise. P values lower than 0.05 were deemed statistically significant.

## 3. Results

### 3.1 The effect of raw force water against chronic exercise fatigue

As shown in Figure 1, after 10 weeks of intensive exercise, the proportion of rats with increasing swimming time in the three exercise groups (Ex CF group, KW-Ex CF group and MW-Ex group) was 44.44%, 88.89% and 37.5%, respectively; and the mean swimming duration of the KW-Ex CF group increased from 326min to 578min (the duration of swimming exercise increased by 252 min), while the other two groups had negative growth (Ex CF group: −11min, MW-Ex group: −8min).

**Figure 1.**
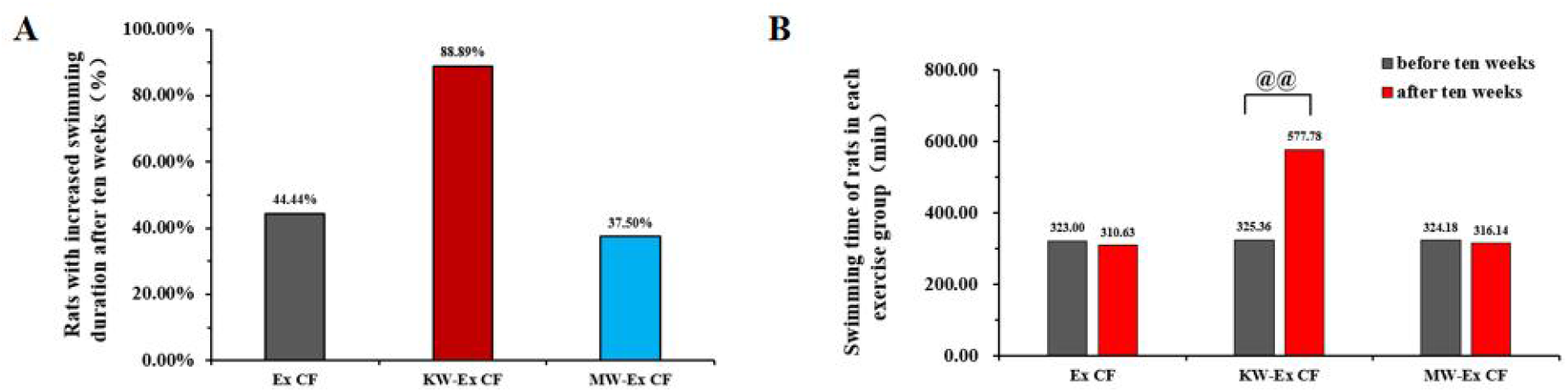
Ultra-small Particle Size Nanobubbles Water (UsNBsW) showed a significantly antifatigue effects in chronic exercise fatigue rats. 1A: Statistics for the proportion of rats in each group (%);1B: Exhaustion exercise time of rats in each group (unit: min). Comparison to ten weeks ago, @@: p<0.01. Ex CF: chronic exercise fatigue model, KW-Ex CF: chronic exercise fatigue model +UsNBsW, MW-Ex: chronic exercise fatigue model +mineral water, each group n=10.

As shown in Figure 2, in Ex CF group, the blood lactate (LA) level (p < 0.05), serum lactate dehydrogenase (LDH) activity (p < 0.05), creatine kinase (CK) activity (p < 0.01), and blood urea nitrogen (BUN) content (p < 0.05) were significantly increased, indicating the successful establishment of the animal model of chronic exercise fatigue. Compared with the Ex CF group of rats, the hemoglobin (Hb) content was significantly increased in the KW-Ex CF group (p < 0.01), serum corticosterone (C) levels (p < 0.05), LA levels (p < 0.01), serum LDH activity (p <0.01), and serum CK activity (p < 0.01) all were significantly decreased. There was no significant difference of serum T / C ratio and BUN levels (p > 0.05); While, when compared to the Ex CF group, the MW-Ex CF group also showed significant changes: such as serum LDH activity (p < 0.01), blood LA level (p < 0.01), and serum CK activity (p < 0.01) were significantly reduced. In addition, the results of this study also showed that the Hb content in KW CF group was significantly increased (p < 0.01), and the Hb content in KW CF group was significantly higher than that in MW-Ex CF group (p < 0.05). It also shows that the Koishio water can significantly improve the Hb content of the body, and has a positive effect on improving chronic exercise fatigue.

**Figure 2.**
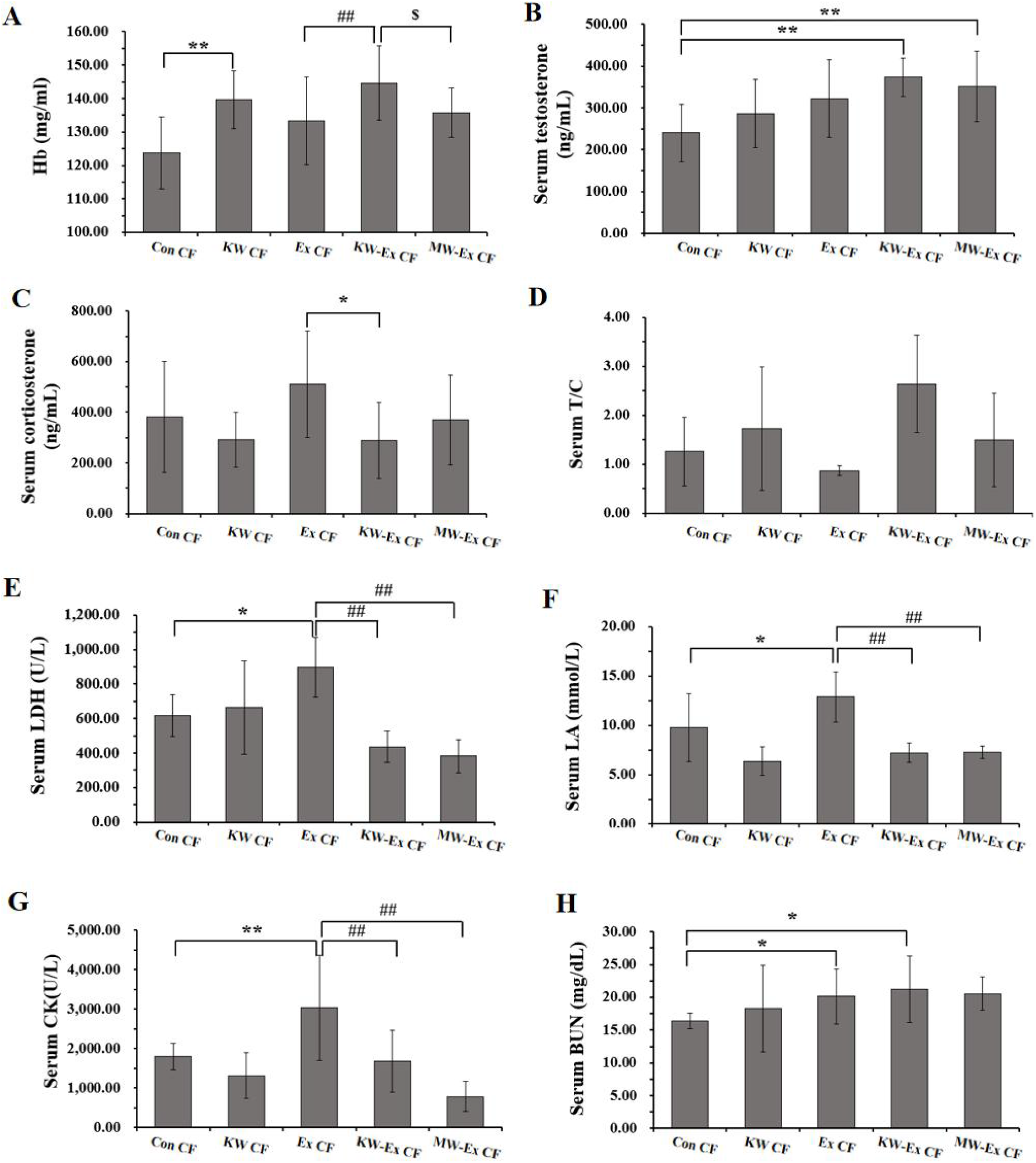
Comparison of anti-fatigue indexes in chronic exercise fatigue rats. 2A: hemoglobin levels in each group (unit: mg/mL); 2B: levels of serum testosterone (T), corticosterone (C) and T/C ratio in each group; 2C: serum lactate dehydrogenase activity in each group (unit: U/LU/L); 2D: blood lactate in each group (unit: U/L);2E: blood urea nitrogen levels in each group (unit: U/Lmg/dL); 2F: serum creatine kinase activity in each group (unit: U/LU/L). Comparison with Con CF, **: p<0.01, *: p<0.05;Comparison with KW-Ex CF group, ##: p<0.01;#: p<0.05;Comparison with KW-Ex CF, $: p<0.05. Con CF: normal feeding; KW-Ex CF: normal feeding + UsNBsW group; Ex CF: Chronic exercise fatigue; KW-Ex CF, Chronic exercise fatigue + UsNBsW group; MW-Ex: Chronic exercise fatigue + mineral water group, each group n=10.

### 3.2 Effect of raw power water on blood glucose and muscle liver glycogen levels in chronic exercise fatigue rats

As shown in Figure 3, there was no significant difference in blood glucose levels after 10 weeks (p > 0.05). Compared with the Ex CF group, muscle glycogen levels in KW-Ex CF group increased but still showed no significant difference (p > 0.05), while liver glycogen levels were significantly higher (p < 0.05) and also significantly higher when compared with the Con CF group (p < 0.05). It suggested that Koishio water does not have a significant effect on blood glucose and muscle glycogen reserve in chronic fatigue rats, but it can significantly increase the liver glycogen level.

**Figure 3.**
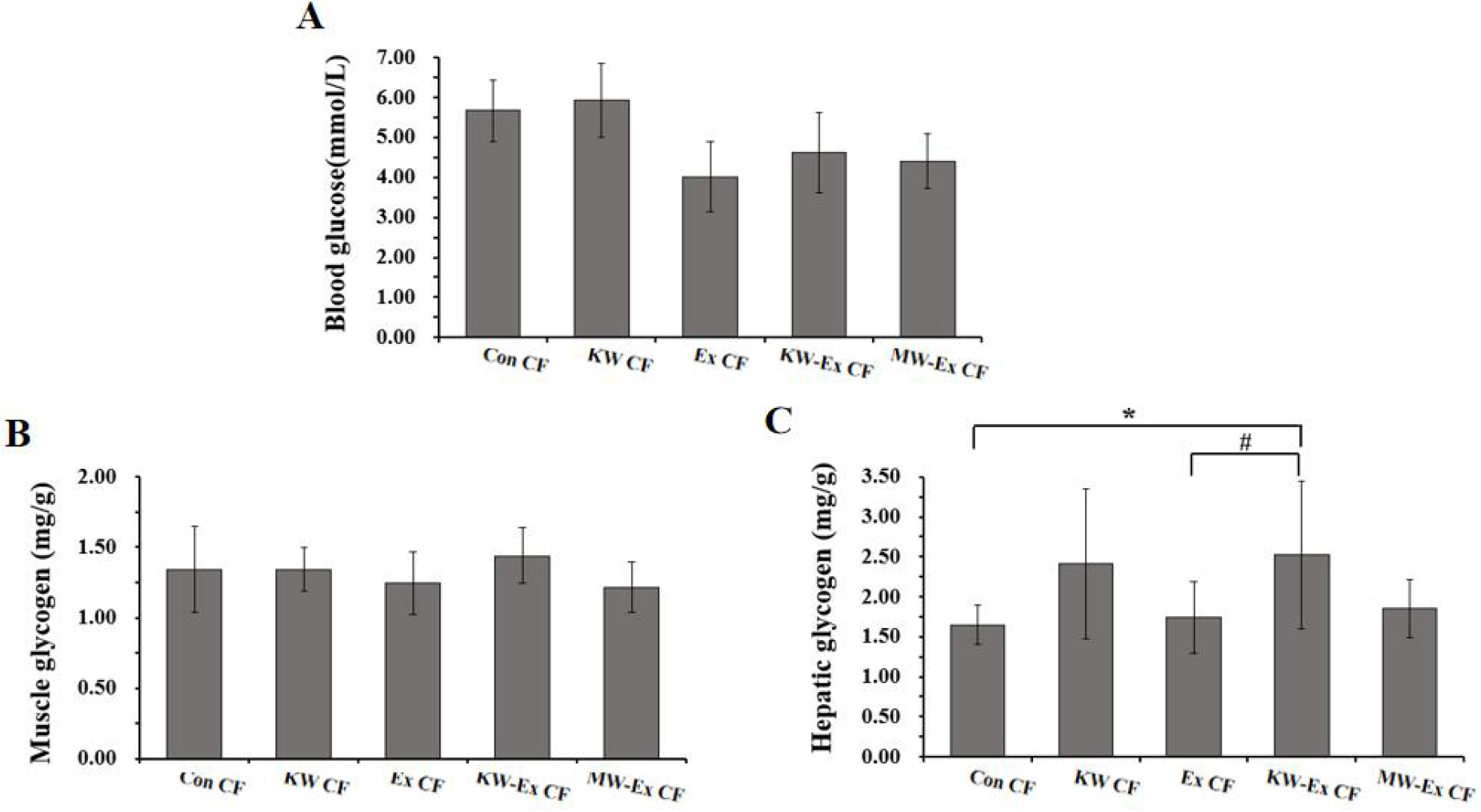
Comparison of blood glucose and muscle hepatic glycogen levels in rats. 3A: Blood glucose levels of the rats in each group (unit: mmol/L); 3B: muscle glycogen levels in each group (unit: mg/g); 3C: hepatic glycogen levelsin each group (unit: mg/g). Comparison with Con CF, *: p<0.05 ; Comparison with KW-Ex CF group, #: p<0.05. Con CF: normal feeding; KW-Ex CF: normal feeding + UsNBsW group; Ex CF: Chronic exercise fatigue; KW-Ex CF, Chronic exercise fatigue + UsNBsW group; MW-Ex: Chronic exercise fatigue + mineral water group, each group n=10.

### 3.3 Effect of stress water on antioxidant indexes in rats with chronic chronic exercise fatigue

As shown in Figure 4, compared with the Con CF group, serum total antioxidant capacity (T-AOC, p < 0.01), superoxide dismutase (SOD, p < 0.01), glutathione peroxidase (GsH-px, p <0.05) were decreased significantly in the Ex CF group, meanwhile, serum malondialdehyde (MDA, p <0.01) and 8-hydroxydeoxyguanosine (8-OH-dG, p < 0.01) were markedly increased; and the skeletal muscle GSH-Px (p < 0.05) decreased significantly and MDA (p < 0.05), 8-OH-dG (p < 0.01) were also significantly increased. When compared with the Ex CF group, the KW-Ex CF group serum T-AOC (p < 0.05), SOD activity (p < 0.01), GSH-Px (p < 0.05) activity increased significantly, the contents of MDA (p < 0.01) and 8-OH-dG (p < 0.05) were decreased significantly, the skeletal muscle MDA (p < 0.01) also show a significantly decreased. However, compared with the Ex CF group, only serum SOD activity (p < 0.05) increased significantly and serum MDA decreased significantly (p < 0.01) in the MW-Ex CF group. These results show that long-term chronic exercise fatigue can cause significant oxidative damage, and the Koishio water has a positive intervention effect on the antioxidant index of long-term chronic exercise fatigue.

**Figure 4.**
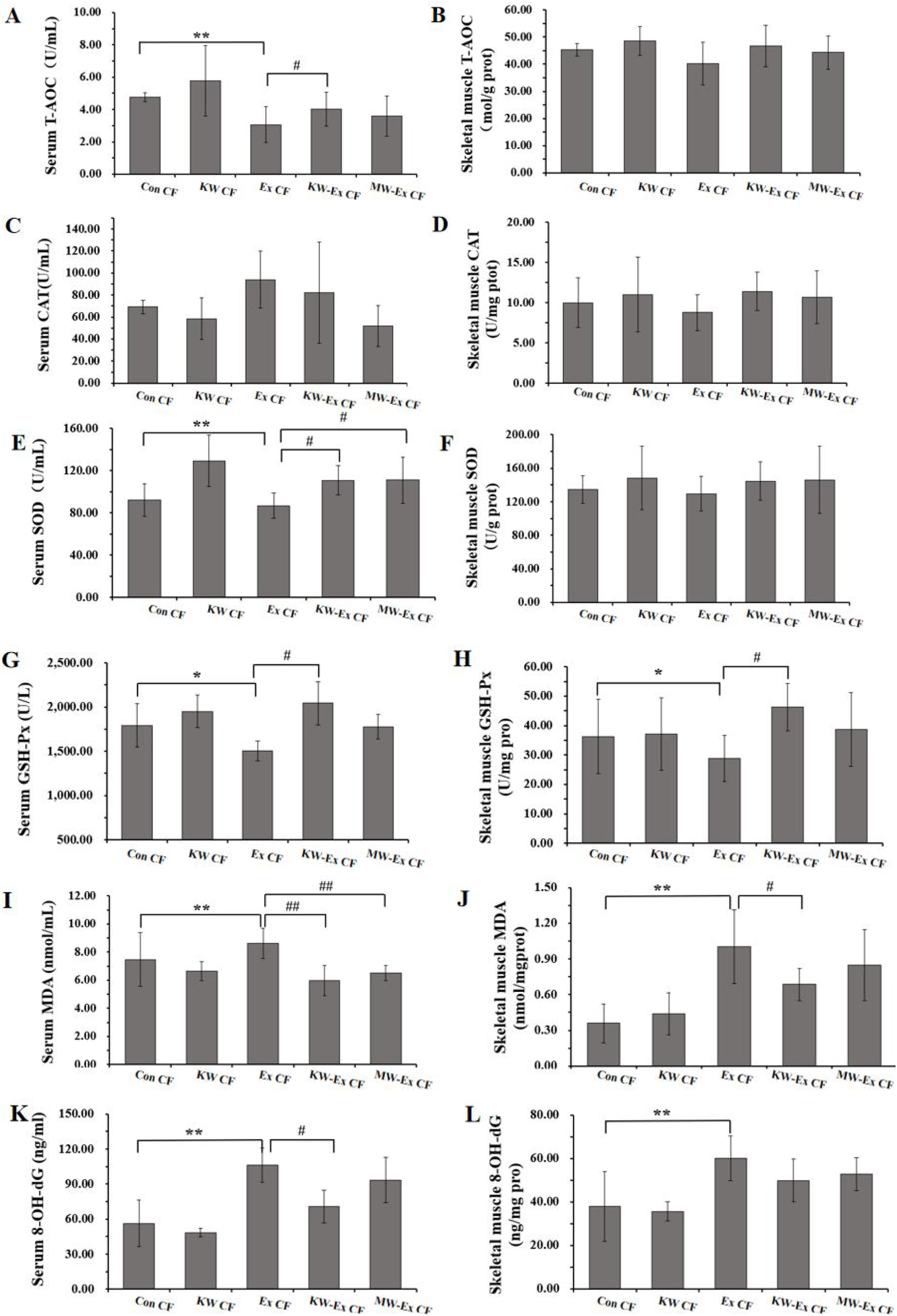
Comparison of antioxidant indexes in rats. 4A-4B: Total antioxidant (T-AOC) levels of serum and skeletal muscle in each group; 4C-4D: serum and skeletal muscle catalase (CAT) activities in each group; 4E-4F: serum and skeletal muscle superoxide dismutase (SOD) activities in each group; 4G-4H: Comparison of serum and skeletal muscle glutathione peroxidase (GSH-Px) levels in each group; 4I-4J: Comparison of serum and skeletal muscle malondialdehyde (MDA) levels in each group of rats; 4K-4L: serum and skeletal muscle 8-hydroxydeoxyguanosine (8-OH-dG) levels in each group. Comparison with Con CF, **: p<0.01, *: p<0.05; Comparison with KW-Ex CF group, ##: p<0.01; #: p<0.05. Con CF: normal feeding; KW-Ex CF: normal feeding + UsNBsW group; Ex CF: Chronic exercise fatigue; KW-Ex CF, Chronic exercise fatigue + UsNBsW group; MW-Ex: Chronic exercise fatigue + mineral water group, each group n=10.

### 3.4 Effect of nuclear force water on C-reactive protein of long-term chronic exercise fatigue rats

As shown in Figure 5, after 10 weeks, serum C-reactive protein (CRP) was not significantly difference (p > 0.05), while in skeletal muscle, CRP level was increased in Ex CF group (p > 0.05); when compared with the Ex CF group, skeletal muscle CRP in KW-Ex CF group (p > 0.05) was obviously decreased. It shows that long-term chronic exercise fatigue can produce inflammatory reaction in the skeletal muscle of rats, and the Koishio water can produce significant improvement effect on C-reactive protein, the inflammatory index of the body after long-term chronic exercise fatigue.

**Figure 5.**
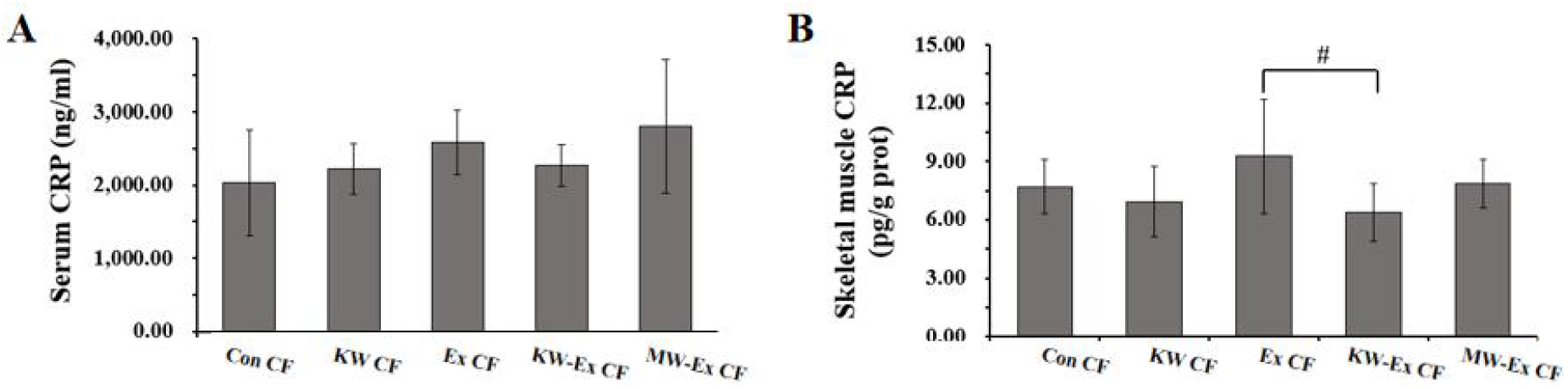
Comparison of inflammatory markers in rats. 5A: serum C-reactive protein (CRP) content in each group; 5B: skeletal muscle CRP content in each group. Comparison with KW-Ex CF group, #: p<0.05. Con CF: normal feeding; KW-Ex CF: normal feeding + UsNBsW group; Ex CF: Chronic exercise fatigue; KW-Ex CF, Chronic exercise fatigue + UsNBsW group; MW-Ex: Chronic exercise fatigue + mineral water group, each group n=10.

### 3.5 Effect of mechanical water on ATP in rats with chronic chronic exercise fatigue

As shown in Figure 6, after 10 weeks, there was no differences of serum ATP between each group (p > 0.05), while in skeletal muscle, ATP content in the Ex CF group (p > 0.05) only show a uptrend (p > 0.05), while the ATP content in KW-Ex CF group was significantly increased (p < 0.01); moreover, the ATP content in skeletal muscle in KW-Ex CF group was significantly higher than that in the MW-Ex CF group (p < 0.05). It suggested that the Koishio water can increase the long-term increase.

**Figure 6.**
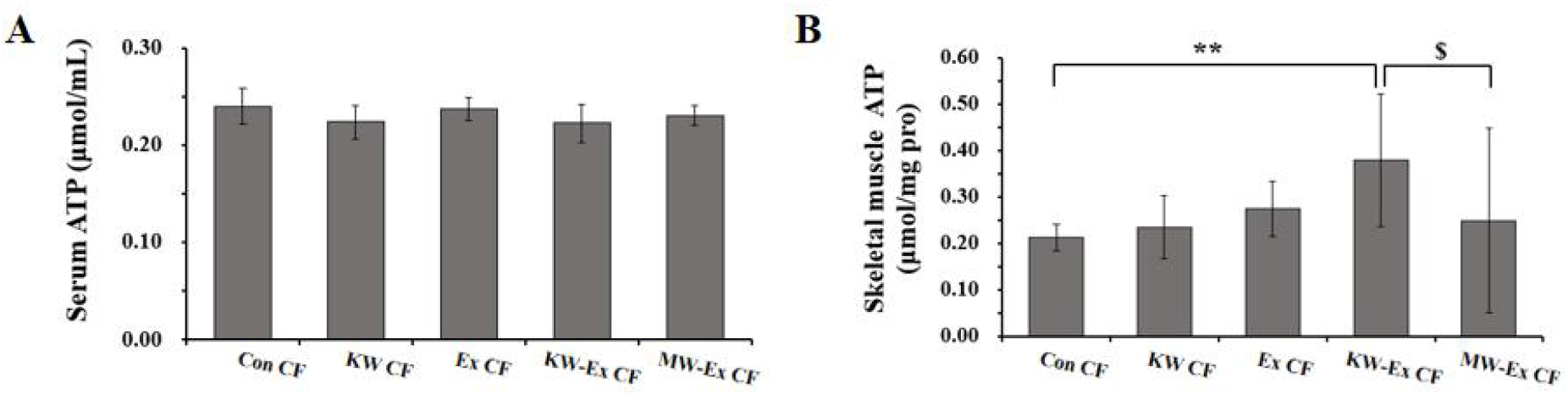
Comparison of adenosine triphosphate levels (ATP) in rats. 6A: serum ATP of the rats in each group; 6B: skeletal muscle ATP of the rats in each group. Comparison with Con CF; **: p<0.01, Comparison with KW-Ex CF, $: p<0.05. Con CF: normal feeding; KW-Ex CF: normal feeding + UsNBsW group; Ex CF: Chronic exercise fatigue; KW-Ex CF, Chronic exercise fatigue + UsNBsW group; MW-Ex: Chronic exercise fatigue + mineral water group, each group n=10.

## 4. Discussions

Chronic fatigue has emerged as a significant health concern, characterized by persistent and distressing sensations of physical and mental exhaustion that can severely impact quality of life and daily functioning. This condition has multifactorial origins, often associated with a wide range of underlying medical conditions, psychological factors, and lifestyle choices. The physiological mechanisms underlying chronic fatigue are complex and involve alterations in metabolic processes, immune system dysregulation, and neurobiological changes. With a prevalence of up to 3% in the general population, chronic fatigue syndrome has garnered attention in both clinical and research settings, highlighting the urgent need for effective therapeutic interventions [8].

In this study, we aimed to investigate the effects of ultra-small particle size nanobubbles water (UsNBsW, namely Koisio water) on physiological and biochemical parameters in a rat model of chronic exercise fatigue. Our research builds upon previous findings that demonstrate the positive impact of exercise on fatigue and overall health, while also exploring the specific role of hydration quality in recovery from exercise-induced fatigue. By systematically analyzing various physiological outcomes, including swimming endurance, biochemical markers of muscle damage, and oxidative stress levels, we provide evidence that UsNBsW may offer a novel approach to mitigating the effects of chronic exercise fatigue. The findings from this study will be discussed in detail, emphasizing their implications for future interventions aimed at enhancing exercise performance and recovery in both clinical and athletic populations [3].

The results of this study provide significant insights into the biological mechanisms through which UsNBsW affects chronic exercise fatigue in rats. The observed increase in hemoglobin levels, coupled with a notable decrease in serum lactate, lactate dehydrogenase, and creatine kinase activity, suggests that KW may enhance oxygen transport and utilization during physical exertion, thereby mitigating the physiological stress associated with chronic fatigue. These findings align with previous literature, which emphasizes the critical role of effective hydration in optimizing athletic performance and recovery through improved metabolic functions and reduced oxidative stress [2]. Moreover, the biochemical alterations suggest that KW may activate specific signaling pathways that promote energy homeostasis and mitochondrial function, ultimately facilitating enhanced physical endurance and recovery [9].

In addition to the biochemical changes observed, the study’s findings on glycogen levels in the liver indicate that KW supports improved energy storage and availability during physical activities. The significant increase in hepatic glycogen levels in the KW group compared to controls suggests a potential mechanism where hydration quality may influence metabolic pathways involved in glycogen synthesis and utilization [10]. This improved energy metabolism is crucial for sustaining prolonged physical activity and could inform dietary and hydration strategies for athletes and individuals engaged in regular exercise. Furthermore, the interplay between ultra-small particle size nanobubbles, glycogen storage, and exercise capacity underscores the importance of tailored nutritional interventions in managing chronic fatigue and enhancing athletic performance [11].

The notable enhancement in antioxidant markers, including superoxide dismutase and glutathione peroxidase activity, alongside reduced levels of lipid peroxidation markers, further illustrates the protective effects of KW against oxidative damage induced by strenuous exercise. This antioxidant response may play a pivotal role in reducing exercise-induced muscle damage and inflammation, aligning with current literature that highlights the importance of antioxidants in mitigating oxidative stress during physical exertion [12]. As such, the findings of this study not only contribute to our understanding of hydration’s role in exercise performance but also propose KW as a potential adjunctive strategy for improving recovery and resilience against chronic exercise fatigue. Future studies are warranted to explore the molecular mechanisms underlying these effects and to evaluate the applicability of KW in clinical and athletic populations [13].

The limitation of this study is the absence of clinical validation limits the applicability of the results to human populations. The study solely relied on animal models, which, while useful, may not fully encapsulate the complexities of human physiology and response to nutritional interventions. Furthermore, the lack of a comprehensive approach incorporating wet lab experiments may hinder a more thorough understanding of the underlying mechanisms. These limitations underscore the need for further research with larger cohorts and clinical trials to validate the efficacy of UsNBsW as a potential intervention for chronic exercise fatigue.

In conclusion, this study reveals the beneficial effects of Koishio water on physiological parameters in rats experiencing chronic exercise fatigue. Notably, improvements in hemoglobin content, liver glycogen levels, and antioxidant capacity were observed, suggesting that such interventions may serve as effective strategies for enhancing recovery and performance in physical activities. These findings provide a foundation for future research and application aimed at exploring nutritional strategies for exercise recovery, with potential implications for practice in sports medicine, military physical fitness rehabilitation and public fitness field.

## References

[1] Nijs J, Almond F, De Becker P, Truijen S, Paul L. Can exercise limits prevent post-exertional malaise in chronic fatigue syndrome? An uncontrolled clinical trial. Clin Rehabil. 2008; 22(5): 426–435. doi:10.1177/0269215507084410

[2] Twomey R, Aboodarda SJ, Kruger R, Culos-Reed SN, Temesi J, Millet GY. Neuromuscular fatigue during exercise: Methodological considerations, etiology and potential role in chronic fatigue. Neurophysiol Clin. 2017; 47(2): 95–110. doi:10.1016/j.neucli.2017.03.002

[3] Oldervoll LM, Kaasa S, Knobel H, Loge JH. Exercise reduces fatigue in chronic fatigued Hodgkins disease survivors--results from a pilot study. Eur J Cancer. 2003; 39(1): 57–63. doi:10.1016/s0959-8049(02)00483-5

[4] Żychowska M. Dapsone: a forgotten and underestimated treatment option for bullous pemphigoid?. Br J Dermatol. 2017; 177(5): 1156–1157. doi:10.1111/bjd.15963

[5] Vybornaya KV, Sokolov AI, Kobelkova IV, Lavrinenko SV, Klochkova SV, Nikityuk DB. Vopr Pitan. 2017; 86(5): 5–10. doi:10.24411/0042-8833-2017-00069

[6] Jammes Y, Stavris C, Charpin C, Rebaudet S, Lagrange G, Retornaz F. Maximal handgrip strength can predict maximal physical performance in patients with chronic fatigue. Clin Biomech (Bristol, Avon). 2020; 73: 162–165. doi:10.1016/j.clinbiomech.2020.01.003

[7] Xu JX, Lin F, Wu YH, Chen ZK, Ma YB, Yang LY. Etiology analysis for term newborns with severe hyperbilirubinemia in eastern Guangdong of China. World J Clin Cases. 2023; 11(11): 2443–2451. doi:10.12998/wjcc.v11.i11.2443

[8] Edmonds M, McGuire H, Price J. Exercise therapy for chronic fatigue syndrome. Cochrane Database Syst Rev. 2004; (3): CD003200. doi:10.1002/14651858.CD003200.pub2

[9] Ma Y, Zheng Y, Zhou Y, Weng N, Zhu Q. Mitophagy involved the biological processes of hormones. Biomed Pharmacother. 2023; 167: 115468. doi:10.1016/j.biopha.2023.115468

[10] Chen TS, Liou SY, Wu HC, et al. New analytical method for investigating the antioxidant power of food extracts on the basis of their electron-donating ability: comparison to the ferric reducing/antioxidant power (FRAP) assay. J Agric Food Chem. 2010; 58(15): 8477–8480. doi:10.1021/jf9044292

[11] Pianosi PT, Emerling E, Mara KC, Weaver AL, Fischer PR. Aerobic fitness in adolescents with chronic pain or chronic fatigue: parallels and mechanisms?. J Rehabil Med. 2017; 49(5): 441–446. doi:10.2340/16501977-2221

[12] Nijs J, Paul L, Wallman K. Chronic fatigue syndrome: an approach combining self-management with graded exercise to avoid exacerbations. J Rehabil Med. 2008; 40(4): 241–247. doi:10.2340/16501977-0185

[13] Wang S, Wang HY, D. J. Discovering causal paths to diabetic nephropathy by combining computable biomedical knowledge with graph mining algorithms. AMIA Annu Symp Proc. 2023; 2022: 1118–1124. Published 2023 Apr 29.

